# Detection of interactions between genetic marker sets and environment in a genome-wide study of hypertension

**DOI:** 10.1101/2023.05.28.542666

**Authors:** Linchuan Shen, Amei Amei, Bowen Liu, Yunqing Liu, Gang Xu, Edwin C. Oh, Zuoheng Wang

## Abstract

As human complex diseases are influenced by the interplay of genes and environment, detecting gene-environment interactions (*G* × *E*) can shed light on biological mechanisms of diseases and play an important role in disease risk prediction. Development of powerful quantitative tools to incorporate *G* × *E* in complex diseases has potential to facilitate the accurate curation and analysis of large genetic epidemiological studies. However, most of existing methods that interrogate *G* × *E* focus on the interaction effects of an environmental factor and genetic variants, exclusively for common or rare variants. In this study, we proposed two tests, MAGEIT_RAN and MAGEIT_FIX, to detect interaction effects of an environmental factor and a set of genetic markers containing both rare and common variants, based on the MinQue for Summary statistics. The genetic main effects in MAGEIT_RAN and MAGEIT_FIX are modeled as random or fixed, respectively. Through simulation studies, we illustrated that both tests had type I error under control and MAGEIT_RAN was overall the most powerful test. We applied MAGEIT to a genome-wide analysis of gene-alcohol interactions on hypertension in the Multi-Ethnic Study of Atherosclerosis. We detected two genes, *CCNDBP1* and *EPB42*, that interact with alcohol usage to influence blood pressure. Pathway analysis identified sixteen significant pathways related to signal transduction and development that were associated with hypertension, and several of them were reported to have an interactive effect with alcohol intake. Our results demonstrated that MAGEIT can detect biologically relevant genes that interact with environmental factors to influence complex traits.

## 1. Introduction

Causes of human complex diseases are multifactorial including the interplay of genes and environment. The effect of environment exposures on disease outcomes can vary across genotypic groups. It has been reported that individuals with certain genetic profiles have elevated disease risk only when they are exposed to an environment in many complex diseases (Lin *and others.*, 2013). For example, many environmental factors such as aging, sex, smoking, diet, stress, air quality and among others influence disease risk, progression and severity (Bhatnagar, 2017, Cosselman, Navas-Acien and Kaufman, 2015). As a result, incorporating gene-environment interactions (*G* × *E*) has become crucial in the study of complex traits. Genome-wide association studies (GWAS) have successfully identified many genetic variants associated with human diseases. However, the estimated effects of these variants are small and only explain small portion of the heritability of complex diseases (Eichler *and others.*, 2010). Several studies have suggested that *G* × *E* may contribute partly to the missing heritability and the detection of *G* × *E* could lead to meaningful implication in fields of public health and personalized medicine (Eichler *and others.*, 2010, Thomas, 2010).

Traditional *G* × *E* analyses focus on evaluating the interactions with genetic variants one at a time (Aschard *and others.*, 2010, Kraft *and others.*, 2007, Manning *and others.*, 2011). Possible limitations in such approaches include the burden of multiple hypothesis testing and lacking consideration of joint effects shared by multiple variants with similar biological functions, resulting in power loss in the analysis (Lin *and others.*, 2013). In recent years, genome-wide search for *G* × *E* has been emerging (Khoury and Wacholder, 2009, Thomas, 2010) and several studies have investigated *G* × *E* from multiple variants in a genetic marker set (Chen, Meigs and Dupuis, 2014, Chi *and others.*, 2021, Jiao *and others.*, 2013, Lin *and others.*, 2019, Lin *and others.*, 2013, Lin *and others.*, 2016, Su, Di and Hsu, 2017, Tzeng *and others.*, 2011, Wang *and others.*, 2020). For a set of common genetic variants, gene-environment set association test (GESAT) was developed using a generalized linear model and ridge regression (Lin *and others.*, 2013). For rare variants, Chen *et al*. proposed INT-FIX and INT-RAN for testing *G* × *E* effect, as well as a joint test, JOINT, that detects the effects of a set of genetic variants as well as their interactions with an environmental factor simultaneously (Chen, Meigs and Dupuis, 2014). They used a beta density function for genetic effect to reflect larger contributions from rare genetic variants. Genetic main effects in their *G* × *E* tests were treated as fixed in INT-FIX or random in INT-RAN, respectively. The three tests were implemented as an R package called rareGE. To assess rare variants by environment interaction, Lin et al. developed the interaction sequence kernel association test (iSKAT) that modeled the main effects of rare variants using weighted ridge regression and allowed the interactions with environment across genetic variants to be correlated (Lin *and others.*, 2016). GESAT, the three tests in the rareGE package and iSKAT are all variance component-based tests that are robust to the signs and magnitudes of the *G* × *E* effects when many variants in a genetic region are non-causal and/or there are mixed beneficial and detrimental variants (Lee, Wu and Lin, 2012, Santorico and Hendricks, 2016, Wu *and others.*, 2011). A unified hierarchical modeling of *G* × *E* effects from a set of rare variants, called mixed effects score test for interaction (MiSTi), which models *G* × *E* effects by a fixed component as well as a random component was developed (Su, Di and Hsu, 2017). They constructed two independent score statistics and combined them using data-adaptive approaches. Simulation studies showed that MiSTi has greater than or comparable power to iSKAT. MiSTi provided a unified regression framework for testing interaction effects between a set of rare variants and an environmental factor where many existing methods can be derived from by constraining certain parameters to be zero. In addition to the above mentioned *G* × *E* tests that were developed under the regression framework, Lin *et al*. proposed a polygenic test of *G* × *E* effect using Bayes factors (Lin *and others.*, 2019). In their adaptive combination of Bayes factors method (ADABF), *G* × *E* effects are assumed to follow a normal distribution. Variants in a genetic region were sorted by Bayes factors and p-values were calculated using a resampling procedure. When there are a few genetic variants interacting with the environmental factor, ADABF had higher power than other methods for detecting *G* × *E* effects.

Complex diseases are influenced by many genetic variants including common and/or rare. Current methods in detecting *G* × *E* mainly focus on the interaction effects of an environmental factor and genetic variants, exclusively for common or rare. Although ADABF considers both common and rare variants in a genetic region, it does not distinguish the effects of the two types of variants in model fitting and hence may overlook the relatively larger contribution from rare variants. Recently, MQS (MinQue for Summary statistics) was developed for estimating variance components in linear mixed models (Zhou, 2017). MQS is based on the method of moments and the minimal norm quadratic unbiased estimation criterion. Compared to the restricted maximum likelihood estimation method (REML), MQS provided unbiased and statistically efficient estimates. It was extended to model the epistatic interactions between genetic variants (Crawford *and others.*, 2017). In this study, we propose two tests to detect interactions between an environmental factor and a set of genetic markers containing both rare and common variants based on the MQS method. We name it as MArginal Gene-Environment Interaction Test with RANdom or FIXed genetic effects (MAGEIT_RAN or MAGEIT_FIX). We assessed the performance of the two tests in detecting *G* × *E* for a set of genetic variants and compared it with existing set-based *G* × *E* methods via simulation studies. Our results demonstrated that both MAGEIT_RAN and MAGEIT_FIX had well controlled type I error. MAGEIT_RAN was most powerful in majority of the simulation scenarios. We applied MAGEIT_RAN and MAGEIT_FIX to a genome-wide analysis of gene-alcohol interaction on hypertension in the Multi-Ethnic Study of Atherosclerosis (MESA) and identified hypertension-related *G* × *E* and pathways.

## 2. Methods

Suppose a phenotype of interest, an environmental variable and genome-wide genetic variants are measured on *n* subjects. Let *y*_*k*_, *E*_*k*_, ***G***_*k*_ = (*G*_*k*1_, *G*_*k*2_, …, *G*_*kp*_)*^T^* and ***X***_*k*_ (*X*_*k*1_, *X*_*k*2_, …, *X*_*km*_)^*T*^ denote the phenotype, environmental variable, genotypes of *p* variants in a genomic region, and *m* non-genetic covariates for the *k*th subject, respectively, for *k* = 1, 2, …, *n*, where *G*_*kj*_ = 0, 1 or 2 depending on whether subject *k* has 0, 1 or 2 copies of minor allele at the *j*th variant. We use ***S***_*k*_ = (*E*_*k*_*G*_*k*1_, *E*_*k*_*G*_*k*2_, …, *E*_*k*_*G*_*kp*_)*^T^* to denote the genetic variants by environment interaction for the *k* th subject. Our goal is to test whether there are interactions between the variant set and environment that influence the phenotype of interest.

### 2.1 Model for continuous phenotype

Let ***y*** = (*y*_1_, *y*_2_, …, *y*_*n*_)^*T*^, ***E*** = (*E*_1_, *E*_2_, …, *E*_*n*_)^*T*^, and ***ε*** = (*ε*_1_, *ε*_2_, ⋯, *ε*_*n*_)^*T*^ denote vectors of the phenotype, environmental variable, and error term of length *n*. We further define an *n* × *m* covariate matrix ***X*** = [***X***_1_, ***X***_2_, ⋯, ***X***_*n*_]^*T*^, an *n* × *p* genotype matrix ***G*** = [***G***_1_, ***G***_2_, ⋯, ***G***_*n*_]^*T*^, and an *n* × *p* matrix ***S*** = [***S***_1_, ***S***_2_, ⋯, ***S***_*n*_]^*T*^ of the *G* × *E* . Then, the following model specifies the relationship between a continuous phenotype ***Y*** and ***X***, ***E***, ***G*** and ***S***

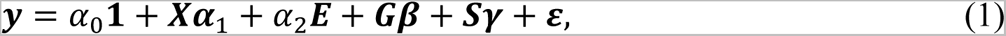

where **1** is an *n* × 1 vector of 1, *α*_0_ is an intercept term, ***α***_1_ = (*α*_11_, *α*_12_, …, *α*_1*m*_)^*T*^, *α*_2_, ***β*** = (*β*_1_, *β*_2_, …, *β*_*p*_)^*T*^ and ***γ*** = (*γ*_1_, *γ*_2_, …, *γ*_*p*_)^*T*^ are regression coefficients for the covariates, environmental factor, genetic variants, and *G* × *E* terms. We further assume that *γ* and ***ε*** follow multivariate normal distributions with 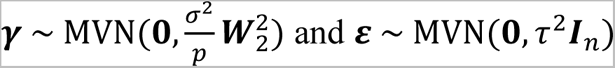, where ***W***_2_ = diag(*w*_21_, *w*_22_, ⋯, *w*_2*p*_) contains weights of the *p G* × *E* terms and ***I***_*n*_ is an identity matrix of dimension *n*.

### 2.2 Marginal gene-environment interaction test

We are interested in testing genetic variants by environment interactions in a genomic region, i.e., testing the null hypothesis *H*_0_: ***γ*** = **0**, which is equivalent to testing *H*_0_: *σ*^2^ = 0. We develop two *G* × *E* tests, in which the genetic main effects ***β*** are modeled as random and fixed, respectively.

When we treat the genetic main effects ***β*** as random, we assume that 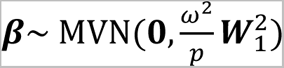, where ***W***_1_ = diag(*w*_11_, *w*_12_, ⋯, *w*_1*p*_) are weights of the *p* variants. We use the MQS method (Zhou, 2017) to estimate the three variance components *ω*^2^, *σ*^2^ and *τ*^2^. In order to eliminate the fix effects *α*_0_, ***α***_1_and *α*_2_in Model (1), we multiply both sides of the model, from left, by a projection matrix ***M***, where ***M*** = ***I*** − ***b***(***b***^*T*^***b***)^−1^***b***^*T*^ with *b* = [***1***, ***X***, *E*]. Then Model (1) becomes

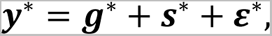

where ***y***^∗^ = ***My***, ***g***^∗^ = ***MGβ*, *s***^∗^ = ***MSγ***, and ***ε*^∗^ = *Mε***. It follows that ***g***^∗^ ∼ MVN(***0***, *ω*^2^***G***^∗^) with 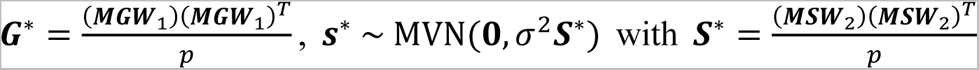, and *ε*^∗^ ∼ MVN(**0**, τ^2^***M***) . Consequently, we have *y*^∗^ ∼ MVN(**0**, *ω*^2^***G***^∗^ + *σ*^2^***S***^∗^ + τ^2^***M***).

We estimate the variance components using the method of moments based on the following set of second moment matching equations,

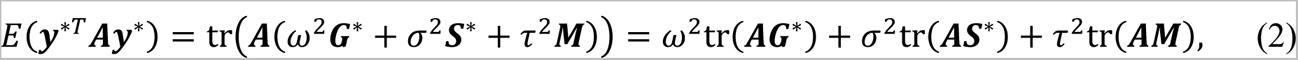

where ***A*** is an arbitrary symmetric non-negative definite matrix (Zhou, 2017). Since there are three unknown parameters (*ω*^2^, *σ*^2^, *τ*^2^), three different ***A***’s are required to obtain parameter estimates.

In the method of moments, the expectation of Eq. (2) is usually replaced with the realized value *y*^∗*T*^***A****y*^∗^. Let ***A***_1_ = ***G***^∗^, ***A***_2_ = ***S***^∗^ and ***A***_3_ = ***M*** (Zhou, 2017), then, the resulting estimates of the variance components are given in a matrix form as

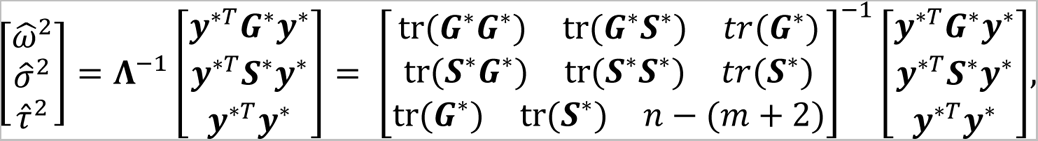

where we used tr(***G***^∗^***M***) = tr(***MG***^∗^) = tr(***G***^∗^), tr(***S***^∗^***M***) = tr(***MS***^∗^) = tr(***S***^∗^), tr(***MM***) = tr(***M***) = *n* − (*m* + 2), and ***y***^∗*T*^***My***^∗^ = ***y***^∗*T*^***y***^∗^ . The variance component estimator 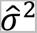 is considered as the test statistic, which we named as MArginal Gene-Environment Interaction Test with RANdom genetic main effects (MAGEIT_RAN). Specifically, the MAGEIT_RAN test statistic is

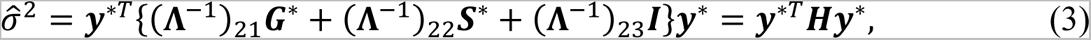

where ***H*** = (**Λ**^−1^)_21_***G***^∗^ + (**Λ**^−1^)_22_***S***^∗^ + (**Λ**^−1^)_23_***I***.

Under *H*_0_: *σ*^2^ = 0, ***y***^∗^ ∼ MVN(**0**, *ω*^2^***G***^∗^ + τ^2^***M***), suggesting that ***y***^∗^ has the same distribution as 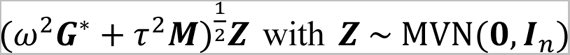. Therefore, the method of moments estimator 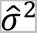 follows the same distribution as 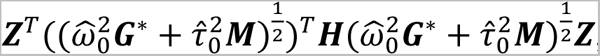, which has a mixture of *χ*^2^ distribution 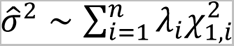. Here, 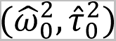 are estimates of (*ω*^2^, *τ*^2^) under the null hypothesis, (*λ*_1_, ⋯, *λ*_*n*_) are eigenvalues of the matrix 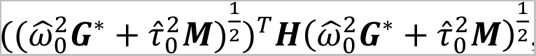, and 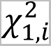 are independent 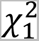 variables (Zhou, 2017).The p-value of 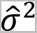 can be evaluated by the Davies method (Davies, 1980, Wu *and others.*, 2011) and Liu-Tang-Zhang approximation (Liu, Tang and Zhang, 2009).

If we treat the genetic main effects ***β*** as fixed, we use the MQS method (Zhou, 2017) to estimate the two variance components *σ*^2^ and *τ*^2^. To eliminate the fix effect terms *α*_0_, ***α***_1_, *α*_2_ and ***β*** in Model (1), we left multiply the model by a projection matrix ***M*** = ***I*** − (***b***^*T*^***b***)^−1^***b***^*T*^with ***b*** = [***1*, *X*, *E*, *G***]. Then the model becomes ***y*^∗^ = *s*^∗^ + *ε*^∗^** and it contains two variance components *σ*^2^ and τ^2^ . Using the method of moments, we obtain the following estimates of the variance components,

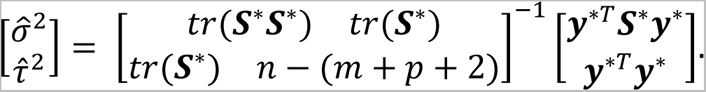

The variance component estimator 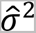 is considered as the test statistic, which we named as MArginal Gene-Environment Interaction Test with FIXed genetic main effects (MAGEIT_FIX). Specifically, the MAGEIT_FIX test statistic is

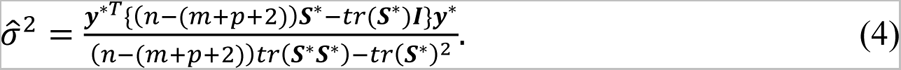

Under 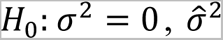 follows a mixture of *χ*^2^ distribution 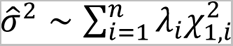 with (*λ*_1_, ⋯, *λ*_*n*_) being the eigenvalues of the matrix 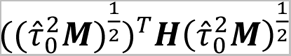.

### 2.3 Model for binary phenotype

We consider a liability threshold model and assume the binary outcome *y*_*k*_ of the *k*th subject is determined by an unobserved continuous liability variable *z*_*k*_, i.e.,

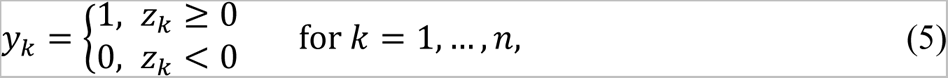

where the underlying liability vector ***z*** = (*z*_1_, *z*_2_, ⋯, *z*_*n*_)^*T*^is specified using Model (1). The full likelihood of the liability threshold mixed effects model is intractable due to an *n*-dimensional integration over the liability variable ***z***. Following the previous studies (Crawford and Zhou, 2018, Engel, Buist and Visscher, 1995, Kuss, Rasmussen and Herbrich, 2005, Tempelman and Gianola, 1993, Williams and Barber, 1998), the liability threshold mixed effects model can be approximated by a linear mixed effects model on 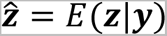, an estimated posterior mean of the liabilities,

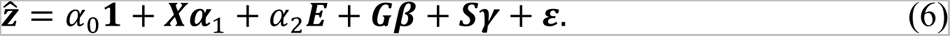

The posterior mean **ẑ** can be obtained by approximation under certain assumptions based on the properties of GWAS data (Crawford and Zhou, 2018). Specifically, we assume that (i) subjects are unrelated, and (ii) both the genetic main effects and interaction effects are small such that the terms ***Gβ*** and ***Sγ*** can be ignored. Under these assumptions, the distribution of the liability variable can be approximated by ***z*** ∼ MVN(*α*_0_**1** + ***Xα***_1_ + *α*_2_*E*, ***I***_*n*_) and **ẑ** is computed as the mean of the following truncated normal distribution (Crawford and Zhou, 2018)

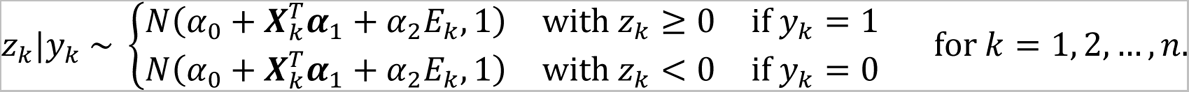

The parameters *α*_0_, ***α***_1_ and *α*_2_ are estimated using a probit model on the phenotype *y*.

To test the interaction effects between a set of genetic variants and an environmental variable on the binary phenotype *y*, we implement MAGEIT_RAN and MAGEIT_FIX on the estimate of the liability variable **ẑ**. To construct MAGEIT_RAN, the liability threshold mixed effects model specified in Eqs (5) and (6) contains three variance components (*ω*^2^, *σ*^2^, τ^2^), where *σ*^2^ represents a measure of interactions between the *p* genetic variants and the environmental variable. In order for the model to be identifiable, we put a constrain on the variance of ***z***, e.g., *ω*^2^ + *σ*^2^ + τ^2^ = 1 (Lee *and others.*, 2011). Similarly, we set *σ*^2^ + τ^2^ = 1 for MAGEIT_FIX.

## 3. Simulation Studies

We conducted simulation studies to evaluate the performance of MAGEIT_RAN and MAGEIT_FIX to detect set-based *G* × *E* effects for both continuous and binary phenotypes, where the variant set contains both common and rare variants. We assessed type I error and empirical power of MAGEIT_RAN and MAGEIT_FIX, and compared them with three set-based *G* × *E* tests, GESAT-W (Lin *and others.*, 2013), aMiSTi (Su, Di and Hsu, 2017), and ADABF (Lin *and others.*, 2019). These three existing methods are popular for *G* × *E* analysis and have well-developed R packages. For fair comparisons, the same weights for rare and common variants were used in all methods except ADABF which does not distinguish common and rare variants and hence no weights were used in the implementation.

### 3.1 Simulation settings

To generate genotypes, we first simulated 100,000 chromosomes over a 5 Kb region using a coalescent model that mimics the linkage disequilibrium (LD) structure and recombination rates of the European population (Schaffner *and others.*, 2005, Shlyakhter, Sabeti and Schaffner, 2014). Then we randomly selected 10 common variants with minor allele frequency (MAF) > 0.05 and 40 rare variants with 0.005 < MAF < 0.05 to compose a set of 50 genetic variants.

We simulated a continuous phenotype using the following trait model,

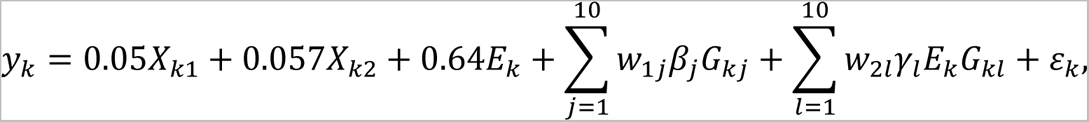

where X_*k*1_ ∼ N(62.4, 11.5^2^) mimicking age and X_*k*2_ ∼ Bernoulli(0.52) mimicking sex (Lin *and others.*, 2013). The 10 genetic variants with main effects and the 10 variants with interaction effects were randomly selected from the set of the 50 variants, independent of *E*. The environmental variable *E* is a Bernoulli random variable taking values of 0 or 1 with a probability of 0.5. The weight of a rare variant in *w*_1*k*_ or *w*_2*l*_ is set to Beta(MAF; 1, 25), the beta density function with parameters 1 and 25 evaluated at the variant’s MAF, and the weight of a common variant in *w*_1*k*_ or *w*_2*l*_is set to *c*Beta(MAF; 0.5, 0.5) with 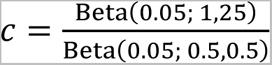 (Ionita-Laza *and others.*, 2013, Madsen and Browning, 2009). The error term *ε*_*k*_ ∼ N(0, 1.5^2^) indicates independent noise.

For a binary trait, we use the following logistic regression model, logit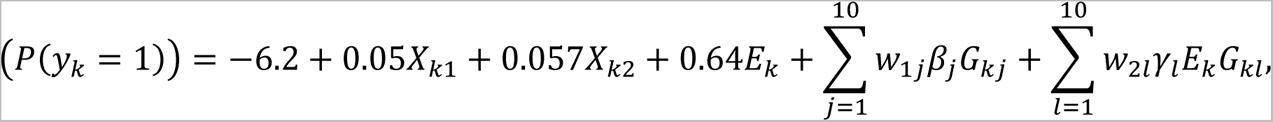

where all parameters are the same as those used in the continuous phenotype model. In all simulation settings, each simulated dataset contains 5,000 subjects (2,500 cases and 2,500 controls for binary phenotype).

In the type I error assessment, we set all *γ*_*l*_ to be 0, i.e., no *G* × *E* effects, and generated 10^6^ datasets containing 50 genetic variants (10 common and 40 rare variants). We considered three scenarios: (1) no genetic main effect, i.e., *β*_*k*_ = 0 for *j* = 1, 2, …, 10; (2) for continuous/binary phenotype, assigning *β*_*k*_ ∼ U(0.07, 0.11) / U(0.08, 0.12) to two randomly selected common variants and *β*_*k*_ ∼ U(0.15, 0.19) / U(0.18, 0.22) to eight randomly selected rare variants; (3) similar to scenario (2) except that half of the common/rare variants have negative effects.

In the power comparison, we designed eight simulation scenarios that differ in three key factors that represent different considerations in the simulation design. The first factor pertains to the presence or absence of genetic main effects; the second factor focuses on the allocation of contributions from common and rare variants; and the third factor considers the direction of genetic main effects and *G* × *E* effects, either all positive effects or half positive and half negative effects. We considered ten variants with *G* × *E* effects, either two common and eight rare variants, or four common and six rare variants. The *G* × *E* effect *γ*_*l*_ was generated from U(0.17, 0.21) and U(0.57, 0.61) for common and rare variants, respectively, for continuous phenotype; and from U(0.28, 0.32) and U(0.86, 0.90) for common and rare variants, respectively, for binary phenotype. The first four simulation scenarios have no genetic main effect and they are as follow: (1) two common and eight rare variants with positive *G* × *E* effects; (2) two common and eight rare variants with *G* × *E* effects, 50% of *γ*_*k*_ > 0 and 50% of *γ*_*k*_ < 0; (3) four common and six rare variants with positive *G* × *E* effects; and (4) four common and six rare variants with *G* × *E* effects, 50% of *γ*_*k*_> 0 and 50% of *γ*_*k*_< 0. The remaining four simulation scenarios have two common and eight rare variants with genetic main effects: (5) *β*_*k*_ was specified the same as in scenario (2) in the type I error assessment, two common and eight rare variants with positive *G* × *E* effects; (6) *β*_*k*_ was specified the same as in scenario (3) in the type I error assessment, two common and eight rare variants with *G* × *E* effects, 50% of *γ*_*k*_ > 0 and 50% of *γ*_*k*_ < 0; (7) *β*_*k*_ was specified the same as in scenario (2) in the type I error assessment, four common and six rare variants with positive *G* × *E* effects; and (8) *β*_*k*_ was specified the same as in scenario (3) in the type I error assessment, four common and six rare variants with *G* × *E* effects, 50% of *γ*_*k*_ > 0 and 50% of *γ*_*k*_ < 0. Power was evaluated using 1,000 simulated datasets in each scenario.

### 3.2 Simulation results

Empirical type I error rate was calculated at the nominal level *α*, for *α* = 0.01, 0.001 and 0.0001, based on 10^6^ replicates, under three simulation scenarios, for both continuous and binary phenotypes (Table 1). In most simulations, the type I error of MAGEIT_FIX was within the 95% confidence interval of the nominal level, while the type I error of MAGEIT_RAN was lower than the nominal level in all simulation settings, especially for binary phenotype, suggesting that the MQS-based testing procedure tends to produce conservative p-values due to the approximation we used to handle binary phenotype (Crawford and Zhou, 2018, Schweiger *and others.*, 2017).

**Table 1.** Empirical type I error of MAGEIT_RAN and MAGEIT_FIX, based on 10^6^ replicates

| Test | Nominal Level | Continuous |  |  | Binary |  |  |
| --- | --- | --- | --- | --- | --- | --- | --- |
|  |  | Scenario 1 | Scenario 2 | Scenario 3 | Scenario 1 | Scenario 2 | Scenario 3 |
| MAGEIT_RAN | 0.01 | **$9.66 \times 10^{-3}$** | **$8.85 \times 10^{-3}$** | **$8.62 \times 10^{-3}$** | **$9.51 \times 10^{-3}$** | **$7.77 \times 10^{-3}$** | **$8.47 \times 10^{-3}$** |
|  | 0.001 | **$8.17 \times 10^{-4}$** | **$6.09 \times 10^{-4}$** | **$5.30 \times 10^{-4}$** | **$7.90 \times 10^{-4}$** | **$4.92 \times 10^{-4}$** | **$5.04 \times 10^{-4}$** |
|  | 0.0001 | **$6.70 \times 10^{-5}$** | **$2.90 \times 10^{-5}$** | **$3.40 \times 10^{-5}$** | **$6.20 \times 10^{-5}$** | **$2.80 \times 10^{-5}$** | **$2.20 \times 10^{-5}$** |
| MAGEIT_FIX | 0.01 | $9.87 \times 10^{-3}$ | $9.98 \times 10^{-3}$ | $1.02 \times 10^{-2}$ | **$9.68 \times 10^{-3}$** | **$9.67 \times 10^{-3}$** | **$9.70 \times 10^{-3}$** |
| | 0.001 | $9.89 \times 10^{-4}$ | $9.99 \times 10^{-4}$ | $9.57 \times 10^{-4}$ | $9.38 \times 10^{-4}$ | $9.58 \times 10^{-4}$ | **$8.99 \times 10^{-4}$** |
| | 0.0001 | $1.01 \times 10^{-4}$ | $9.80 \times 10^{-5}$ | $8.80 \times 10^{-5}$ | $9.00 \times 10^{-5}$ | $8.70 \times 10^{-5}$ | $9.20 \times 10^{-5}$ |

The 95% confidence interval of a nominal level $\alpha$ was calculated as $\alpha \pm 1.96\sqrt{\alpha(1-\alpha)/10^6}$ . Specifically, the 95% confidence intervals are $(9.80 \times 10^{-3}, 1.02 \times 10^{-2})$ for $\alpha = 0.01$ , $(9.38 \times 10^{-4}, 1.06 \times 10^{-3})$ for $\alpha = 0.001$ , and $(8.04 \times 10^{-5}, 1.20 \times 10^{-4})$ for $\alpha = 0.0001$ . Rates outside of the 95% confidence interval are in bold.

Empirical power was calculated at the significant level of 10^−4^, based on 1,000 simulation replicates. Figures 1 and 2 demonstrate the power results of the five methods, MAGEIT_RAN, MAGEIT_FIX, GESAT-W, aMiSTi and ADABF, under eight simulation scenarios, for continuous and binary phenotypes, respectively. MAGEIT_RAN had comparable to higher power than the other methods across all simulation scenarios. We observed similar patterns for continuous and binary phenotypes. MAGEIT_RAN was much more powerful than other tests when there was no genetic main effect (Scenarios 1-4). For continuous traits, MAGEIT_FIX had comparable power to GESAT-W and higher power than aMiSTi in all simulation scenarios. For binary phenotypes, GESAT-W was comparable or more powerful than MAGEIT_FIX and ADABF. When the *G* × *E* effects had mixed positive and negative directions (Scenarios 2,4,6,8), aMiSTi had the lowest power for both continuous and binary phenotypes. Since aMiSTi is a combination of burden and variance component test, it loses power when there are both protective and detrimental variants in the genomic region being tested (Basu and Pan, 2011).

**Figure 1.**
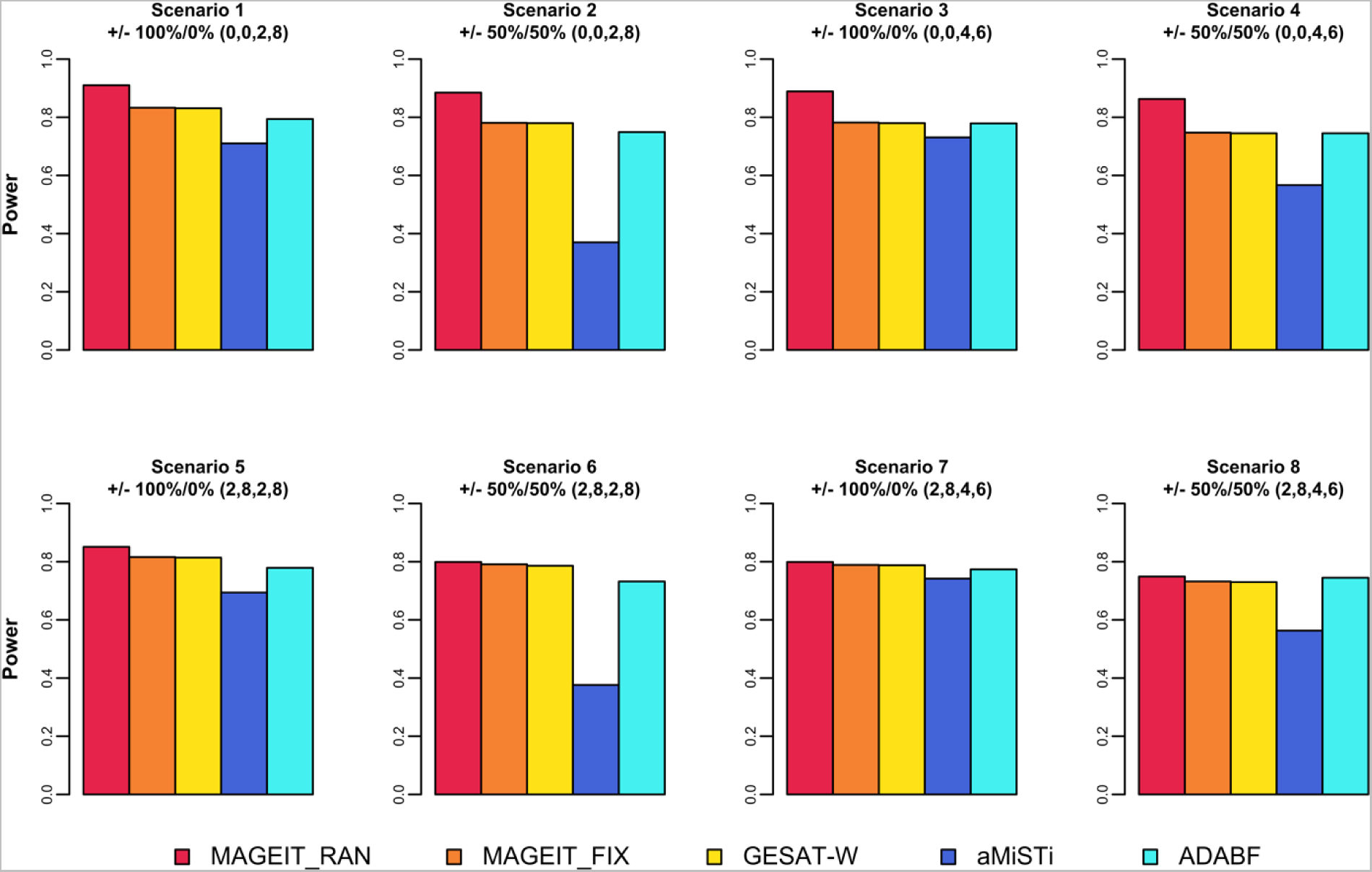
Empirical power of MAGIT_RAN, MAGIT_FIX, GESAT-W, aMiSTi and ADABF for a continuous phenotype.

**Figure 2.**
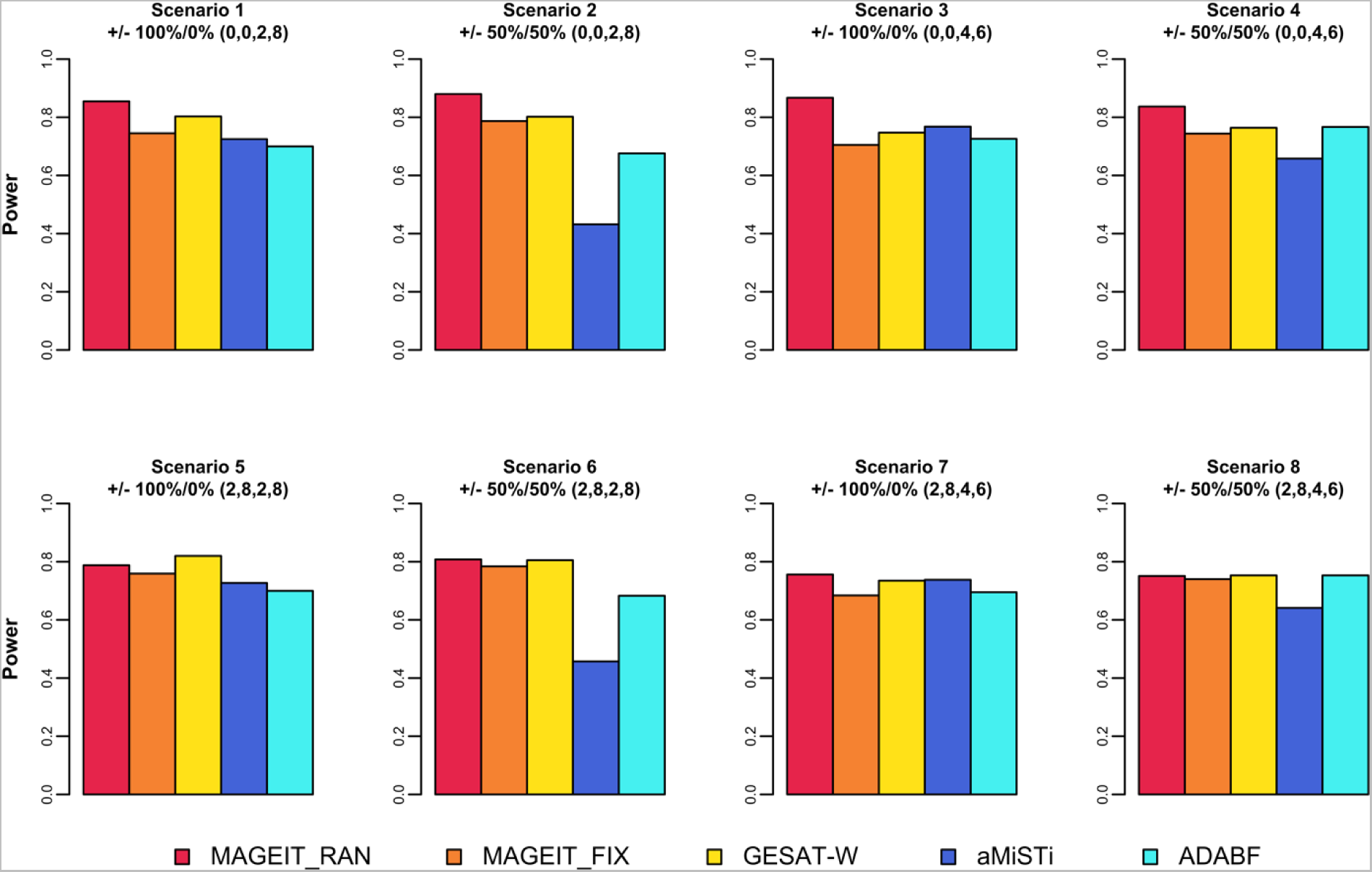
Empirical power of MAGIT_RAN, MAGIT_FIX, GESAT-W, aMiSTi and ADABF for a binary phenotype.

## 4. Application to MESA data

To demonstrate the utility of our proposed methods, we performed a genome-wide analysis of gene-alcohol interaction on hypertension in MESA (Bild *and others.*, 2002). MESA is a large longitudinal study of subclinical cardiovascular diseases including more than 6,800 participants. We analyzed the hypertension outcome measured at the first physical examination of 6,403 participants, consisting of 2,851 subjects with hypertension and 3,552 subjects without hypertension. The participants cover a diverse group of subjects including white (39.3%), African American (26.1%), Hispanic (22.5%), and Asian (12.1%). Alcohol usage (consumption of alcoholic beverages currently or formerly) was treated as an environmental variable, with 6,379 responses including 5,058 YESs and 1,321 NOs.

Samples were genotyped using the Affymetrix Human SNP Array 6.0. After data cleaning, IMPUTE2 (Howie, Donnelly and Marchini, 2009) was used for imputation with the 1000 Genome Phase 3 data as a reference panel. We excluded subjects whose proportion of successfully imputed variants < 5% or empirical inbreeding coefficients > 0.05, resulting in 6,424 subjects for further analysis. The following quality-control criteria were applied: (1) call rate > 95%, (2) MAF > 0.5%, and (3) Hardy-Weinberg *χ*^2^ statistic p-value > 10^−6^, resulting in a final set of 8,540,864 variants. In the gene-based *G* × *E* analysis, we restricted analysis on protein-coding regions based on the reference genome GRCh37 (Frankish *and others.*, 2019). In total, there were 18,977 genes on the 22 chromosomes and the number of variants in each gene region ranges from 2 to 5000, with a medium number of 383. Upon integrating the hypertension, alcohol usage and genotype data, a final set of 6,375 individuals are retained for downstream analyses.

### 4.1 Analysis of *G* × *E* effect

We performed genome-wide tests of gene-alcohol interaction effects on hypertension using all five methods, MAGEIT_RAN, MAGEIT_FIX, GESAT-W, aMiSTi, and ADABF. Age at the first exam, sex, and the top ten principal components (PCs) of the genetic relationship matrix were included in the analysis. The top ten PCs were calculated using the LD pruned variants with MAF > 0.05 to control for population structure.

MAGEIT_RAN and aMiSTi showed no evidence of inflation, with the genomic control inflation factors of 0.966 and 0.997, respectively. The *G* × *E* test assuming fixed genetic main effects, MAGEIT_FIX, and the Bayes factor-based test, ADABF, were conservative, with the genomic control inflation factors of 0.822 and 0.826, respectively. The genomic control inflation factor was 1.403 for GESAT-W. Therefore, we further adjusted the results of GESAT-W using genomic control.

No genes reached genome-wide significance at the p-value threshold of 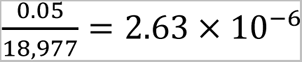, commonly-used in gene-based analyses (Epstein *and others.*, 2015). Table 2 lists the top genes for which at least one of the five tests gives a p-value < 10^−4^. The gene *CCNDBP1* had the smallest p-value, detected by MAGEIT_RAN (p-value = 2.80 × 10^−5^) at a significance level of 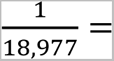 5.27 × 10^-5^, a suggestive significance threshold in genome-wide scan (Lander and Kruglyak, 1995). The p-value of *EPB42* (p-value = 5.98 × 10^−5^) is close to the suggestive significance threshold, generated by MAGEIT_RAN. Both *CCNDBP1* and *EPB42* are located at 15q15.2. The cytogenetic region 15q15 has previously been reported to be associated with blood pressure (Kraja *and others.*, 2005). Moreover, *EPB42* was shown to be significantly down-regulated in heavy drinkers after exposed to psychological stress (Beech *and others.*, 2014, Chen *and others.*, 2021, Ma *and others.*, 2022).

**Table 2.** Genes with p-value < 10^−4^ in at least one of the tests in the MESA data

| Chr | Gene | # SNP | Region | MAGEIT_RAN | MAGEIT_FIX | GESAT-W | aMiSTi | ADABF |
| --- | --- | --- | --- | --- | --- | --- | --- | --- |
| 15 | *CCNDBP1* | 237 | 15q15.2 | **$2.80 \times 10^{-5}$** | $2.03 \times 10^{-3}$ | $6.28 \times 10^{-3}$ | $3.32 \times 10^{-2}$ | $4.90 \times 10^{-2}$ |
| | *EPB42* | 269 | 15q15.2 | **$5.98 \times 10^{-5}$** | $2.05 \times 10^{-3}$ | $1.12 \times 10^{-2}$ | $3.97 \times 10^{-2}$ | $1.00 \times 10^{-1}$ |

The smallest p-values among the five tests at the given genes are in bold.

### 4.2 Pathway analysis

Functional pathway analysis was conducted on genes that had *G* × *E* to identify enriched pathways related to hypertension, using MetaCore^TM^. The top genes for which at least one of the five tests had a p-value < 5 × 10^−3^ were selected. Fisher’s exact test was used to determine whether the gene list was enriched for a functional pathway. At the false discovery rate (FDR) < 0.05, there are 16 significant pathways that were reported to be related to hypertension (Table 3). Of particular interest are eight signaling pathways related to development and signal transduction that are relevant to hypertension and alcohol drinking. The first three pathways include a signal transduction pathway related to ERK1/2 signaling (p-value = 9.66 × 10^−5^, FDR = 1.08 × 10^−2^) and two development pathways related to activation of ERK by alpha-1 adrenergic receptors (p-value = 4.82 × 10^−3^, FDR = 4.92 × 10^−2^) and EPO-induced MAPK (p-value = 4.82 × 10^−3^, FDR = 4.92 × 10^−2^). Mitogen-activated protein kinases (MAPKs) are a group of serine/threonine kinases that include extracellular signal-regulated kinase 1/2 (ERK1/2), c-Jun N-terminal kinases (JNK1/2/3), and p38 (El-Mas and Abdel-Rahman, 2019). Highly conserved in eukaryotes, the MAPK signaling pathway has been implicated in cardiac remodeling and myocardial damage (Liu and Molkentin, 2016). For example, cardiac-specific ERK1/2 knockout mice developed cardiac dilation and eccentric growth of the heart (Kehat *and others.*, 2011), suggesting that ERK1/2 can regulate critical signal transduction pathways. In addition, p38 family members can participate in both protective and deleterious actions in the stress myocardium, demonstrating a key role for MAPKs proteins in cardiac physiology (Romero-Becerra *and others.*, 2020). The fourth pathway is a developmental module related to vascular endothelial growth factor (VEGA) signaling and activation (p-value = 3.97 × 10^−3^, FDR = 4.50 × 10^−2^). The VEGF signaling pathway plays a vital role in the vasculogenesis and angiogenesis in both embryo and adult (Zachary and Gliki, 2001). During neovascularization, VEGF is involved in gene expression, vascular permeability, and the migration, proliferation, and survival of cells (Shibuya, 2011). Studies indicated that VEGF blockade leads to endothelial dysfunction and the inhibition of VEGF-dependent vasodilatory pathways (Robinson *and others.*, 2010). These mechanisms together with the loss of microvascular capillary density through capillary rarefaction cause systemic vasoconstriction and hence resulting in hypertension (Robinson *and others.*, 2010). An animal study also suggested that physiologically relevant levels of alcohol consumption may associated with the stimulation of VEGF expression and angiogenesis (Tan *and others.*, 2007). The fifth one is a development pathway related to positive regulation of WNT/Beta-catenin signaling in the nucleus (p-value = 1.87 × 10^−3^, FDR = 3.92 × 10^−2^). WNT/Beta-catenin signaling pathway, also called canonical WNT signaling pathway in this context, is active in adult cardiac tissue after many cardiac injuries (Ozhan and Weidinger, 2015). WNT/Beta-catenin governs several elements of the renin-angiotensin system (RAS) containing angiotensinogen, renin, angiotensin-converting enzyme, and AT1 receptor (L Ruby *and others.*, 2010, Xiao *and others.*, 2019). Animal studies have shown that WNT2-deficient (another canonical Wnt signaling component) mice exhibit vascular abnormalities, abnormal vascular patterns, and increased vascular fragility (Cattelino *and others.*, 2003, Iso *and others.*, 2006). Interestingly, chronic ethanol consumption affects the activation of WNT/Beta-catenin signaling pathway and WNT/Beta-catenin directly controls alcohol-induced big potassium (BK) internalization (Mercer, Hennings and Ronis, 2015, Velázquez-Marrero *and others.*, 2016). The sixth one is a development pathway related to WNT and Notch signaling in early cardiac myogenesis (p-value = 2.35 × 10^−3^, FDR = 4.00 × 10^−2^). The Notch-mediated signaling, together with other signaling pathway such as VEGF and WNT, play a crucial role in vascular development and angiogenesis (Caliceti *and others.*, 2014). Alternations of Notch signaling is responsible for abnormal blood vessel and heart malformations (Zhou and Liu, 2014). Studies shown that in human coronary artery endothelial cells, ethanol activates notch pathway (Morrow *and others.*, 2010, Morrow *and others.*, 2014). The seventh pathway is a signal transduction pathway related to angiotensin II signaling via beta-arrestin (p-value = 9.17 × 10^−4^, FDR = 3.00 × 10^−2^). Angiotensin II (Ang II), a potent vasoconstrictor and a major effector molecule of renin-angiotensin system (RAS), participates in atherosclerosis and cardiovascular remodeling and raises blood pressure by exploiting various signaling cascades like WNT/beta-catenin (Benigni, Cassis and Remuzzi, 2010, Forrester *and others.*, 2018, Fyhrquist, Metsärinne and Tikkanen, 1995, Kawai *and others.*, 2017, Zhou and Liu, 2016). Animal studies reveal that angiotensinogen genes affect alcohol drinking behavior through Ang II (Fitts, 1993, Maul *and others.*, 2001). The last pathway is a signal transduction pathway related to adenosine A1 receptor signaling (p-value= 1.06 × 10^−4^, FDR= 1.08 × 10^−2^). Adenosine modulates cardiovascular function and produces bradycardia and hypotension when mediated systematically (Barraco *and others.*, 1987, Evoniuk, von Borstel and Wurtman, 1987). Activation of adenosine A1 receptor causes contraction of vascular smooth muscle and the adenosine A1 receptor agonists produce decreases in blood pressure and heart rate (Schindler *and others.*, 2005). It has been observed that raised adenosine levels mediate the ataxic and sedative/hypnotic effects of ethanol through activation of A1 receptors in the cerebellum, striatum, and cerebral cortex (L Ruby *and others.*, 2010). A1 agonists have been shown to decrease anxiety-like behavior, tremor, and seizures during acute ethanol withdrawal in mice (Kaplan *and others.*, 1999).

**Table 3.** Pathways with FDR < 0.05 in the MESA data

| Pathway | P-value | FDR |
| --- | --- | --- |
| Signal transduction_ERK1/2 signaling pathway | $9.66 \times 10^{-5}$ | $1.08 \times 10^{-2}$ |
| Signal transduction_Adenosine A1 receptor signaling pathway | $1.06 \times 10^{-4}$ | $1.08 \times 10^{-2}$ |
| Signal transduction_Adenosine A3 receptor signaling pathway | $4.76 \times 10^{-4}$ | $2.16 \times 10^{-2}$ |
| Development_Thromboxane A2 signaling pathway | $5.57 \times 10^{-4}$ | $2.17 \times 10^{-2}$ |
| Translation_Translation regulation by Alpha-1 adrenergic receptors | $6.96 \times 10^{-4}$ | $2.47 \times 10^{-2}$ |
| Signal transduction_S1P2 receptor inhibitory signaling | $8.58 \times 10^{-4}$ | $2.92 \times 10^{-2}$ |
| Signal transduction_Angiotensin II signaling via Beta-arrestin | $9.17 \times 10^{-4}$ | $3.00 \times 10^{-2}$ |
| Regulation of CFTR activity (normal and CF) | $1.26 \times 10^{-3}$ | $3.21 \times 10^{-2}$ |
| Development_Positive regulation of WNT/Beta-catenin signaling in the nucleus | $1.87 \times 10^{-3}$ | $3.92 \times 10^{-2}$ |
| Signal transduction_S1P1 receptor signaling | $2.13 \times 10^{-3}$ | $3.92 \times 10^{-2}$ |
| Development_WNT and Notch signaling in early cardiac myogenesis | $2.35 \times 10^{-3}$ | $4.00 \times 10^{-2}$ |
| Nociception_Nociceptin receptor signaling | $2.67 \times 10^{-3}$ | $4.19 \times 10^{-2}$ |
| Development_VEGF signaling and activation | $3.97 \times 10^{-3}$ | $4.50 \times 10^{-2}$ |
| Development_Estrogen-independent activation of ESR1 and ESR2 | $4.53 \times 10^{-3}$ | $4.80 \times 10^{-2}$ |
| Development_Activation of ERK by Alpha-1 adrenergic receptors | $4.82 \times 10^{-3}$ | $4.92 \times 10^{-2}$ |
| Development_EPO-induced MAPK pathway | $4.82 \times 10^{-3}$ | $4.92 \times 10^{-2}$ |

## 5. Conclusion

Human complex diseases are influenced by both genetic variation and interactions between gene and environmental factors. Many disease-associated genes have already been identified. Consequently, detecting and understanding gene-environment interactions becomes an important task for disease risk prediction (Hunter, 2005). In this study, we developed two methods MAGEIT_RAN and MAGEIT_FIX to detect the interaction between an environmental factor and gene sets where the genetic main effects were modeled as random or fixed, respectively. Both tests can be applied to continuous and binary phenotypes. Our methods is based on the MQS estimation (Zhou, 2017), which has been applied in MAPIT (Marginal ePIstasis Test) (Crawford *and others.*, 2017) and LT-MAPIT (liability threshold marginal epistasis test) (Crawford and Zhou, 2018) to detect gene-gene interactions. Our methods not only apply the MQS estimation to detect gene-environment interaction but also extends their methods by modeling genetic main effects as random in MAGEIT_RAN. Since variants in a genomic region can be either protective or deleterious and their effect sizes may vary, modeling genetic effects as random, as in MAGEIT_RAN, can capture different directions and magnitude of the genetic effects.

We compared the performance of MAGEIT_RAN and MAGEIT_FIX and three set-based *G* × *E* tests through simulations and real data analysis. In the simulation study, we demonstrated that MAGEIT_FIX had well-controlled type I error rate while MAGEIT_RAN was slightly conservative, especially for binary phenotypes, due to approximations used when specifying MAGEIT on binary phenotypes. MAGEIT_RAN was overall the most powerful among the five methods across all simulation settings. Application of MAGEIT_RAN and MAGEIT_FIX to the MESA hypertension data identified two genes, *CCNDBP1* and *EPB42*, located at the cytogenetic region 15q15.2 which has been reported to be associated with blood pressure. The *EPB42* gene was reported to be significantly down-regulated in heavy drinkers after exposed to psychological stress. Moreover, we identified 16 significant pathways that were related to hypertension, among them eight signaling transduction and development pathways are related to hypertension and alcohol usage. Given the established role of the genes and pathways we identified, MAGEIT has been shown to be able to detect biologically relevant genes that interact with environmental factors to influence complex traits.

There are several limitations in our methods. First, the type I error of MAGIT_RAN is conservative, particularly when there are genetic main effects, thus resulted in slight power loss in simulation scenarios 5-8. This is because the variation of the estimated variance component 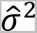 is larger when there are genetic main effects compared with no genetic main effect. Additionally, in MAGEIT_RAN, the regression coefficients *β_j_* of the genetic main effects are assumed to be independent and *γ*_j_ of the *G* × *E* interactions as well. In reality, it is possible there exist correlations among these effects in a genomic region. This assumption may contribute to power loss, particularly in cases where most variants interact with the environmental factor and the effects of interactions are in the same direction. Nevertheless, considering the inherent complexities of linkage disequilibrium and haplotype effects, it is more appropriate to consider potential correlations among these coefficients. Given this, our model can be expanded to accommodate correlations among variants in a genomic region.

## 6. Software

Code to reproduce the results of the article is available at https://github.com/ZWang-Lab/MAGEIT.

